# ROSMAP-Compass: a data-harmonised, AI-ready atlas of 22 million single nuclei from the ROSMAP cohort

**DOI:** 10.1101/2025.08.11.668964

**Authors:** Matthias Flotho, Ian Ferenc Diks, Philipp Flotho, Pascal Hirsch, Viktoria Wagner, Friederike Grandke, Andreas Keller

## Abstract

The Religious Orders Study and Memory and Aging Project (ROSMAP) cohort has generated the world’s most comprehensive single-cell transcriptomic resource for Alzheimer’s disease research. Naturally, in a project spanning multiple years with dozens of research groups involved, the resulting data landscape shows fragmentation across sequencing chemistries, protocols, and pipelines. This presents both a challenge and a unique opportunity for harmonized, collaborative analysis. Following an early data integration strategy and complete realignment of all single nucleus RNA sequencing data, we generated a fully harmonized resource: ROSMAP-Compass, comprising more than 22 million high-quality nuclei from 2,058 donors in multiple brain regions from the ROSMAP and Neuro Psychiatric Symptoms (NPS-AD) cohorts. Through systematic curation and unified reprocessing, we addressed substantial technical challenges including chemistry-specific biases, cross-study batch effects, and sample redundancies across multiple studies spanning different time periods and research groups. ROSMAP-Compass demonstrates the critical importance of systematic data harmonization when integrating large-scale single-cell datasets from multiple sources. By combining open science principles with cutting-edge AI integration, we provide both a critical resource for understanding Alzheimer’s disease heterogeneity and a blueprint for making complex biomedical data accessible to the global research community. The full resource, interactive web portal, and LLM compatible API are freely available, empowering researchers worldwide to accelerate discovery in neurodegenerative diseases.

## 1 Introduction

The Religious Orders Study and Memory Aging Project (ROSMAP)[3] cohort established by the Rush Alzheimer’s Disease Center (RADC), represents one of the most comprehensive longitudinal studies of aging and Alzheimer’s disease. With more than **3,500** participants enrolled, including more than 1,300 deceased donors, this cohort provides unparalleled richness comparable to the Allen Brain Atlas (Figure 1a). The Study captures multiple data modalities from participants, including small non-coding RNA profiles [20], imaging data, and extensive clinical parameters. ROSMAP distinguish in capturing the full spectrum of Alzheimer’s disease pathology. Analysis of deceased patients (N=1,596) reveals complex relationships between neuropathological assessments and clinical diagnoses (Figure 1b). Notably, CERAD scores, which quantify neuritic plaque density, show variable correlation with cognitive diagnoses, highlighting the heterogeneity of AD presentation and the importance of molecular profiling to understand disease mechanisms. The single-cell era of ROSMAP began in 2019 when Mathys et al. first characterized **48** patients using 10x Chromium v2 technology [16] (Figure 1c). This seminal study employed rigorous donor selection criteria and demonstrated strong correlation between clinically diagnosed and pathological confirmed Alzheimer’s disease. Since this foundational work, multiple large-scale studies have expanded our understanding of AD at single-cell resolution (Figure 1b,c). Key contributions include Blanchard et al., who sequenced 24 additional patients to investigate APOE-related mechanisms (Figure 1c). The Tsai laboratory has continued this work with the Mathys et al. (2023) study [17], currently in press at Cell, and most recently published a multi-region brain atlas in 2024 [15]. This multiregional approach has provided crucial insights into the spatial heterogeneity of AD pathology. Parallel efforts from the De Jager laboratory have further enriched the ROSMAP single-cell landscape. In 2024, Fujita et al.[9] and Green et al.[11] published a comprehensive multimodal atlas of the ROSMAP cohort (Figure 1c). Additionally, Lee et al. [14] incorporated data from the Neuro Psychiatric Symptoms in AD (NPS-AD) cohort, which provides comparable metadata quality and expands the biological diversity of samples analyzed (Table 1). Although each of these studies represents a significant advance in understanding the molecular basis of AD, the distributed nature of the data presents challenges to the broader research community. Non-trivial cell overlaps between studies, combined with technical variations introduced by different sequencing chemistries, complicate direct integration and comparison of results. This uniquely rich cohort, with its many sequencing studies spanning different technologies, time points, and research groups, calls for a unified and accessible resource that enables cross-study integration and coherent downstream analysis, leveraging the resource for novel discoveries in AD. We thus developed a curated and harmonized resource integrating all available ROSMAP single-cell RNA seq data. Given the complexity and diversity of the available singlecell datasets, one key design decision was the stage at which to integrate data. In principle, integration can occur at multiple levels, ranging from raw reads (early), to count matrices (intermediate), to aggregated gene-level summaries (late). Each approach has distinct advantages and limitations. We chose an early integration strategy, aligning all available reads using a unified reference and consistent pipeline. This allowed us to minimize technical artifacts introduced by differing chemistries and processing methods, and to maximize comparability across donors and studies. Our goal is to make this invaluable dataset broadly accessible to the open science community, enabling researchers worldwide to leverage the full richness of the ROSMAP cohort and build upon the foundational work of these pioneering studies.

**Table 1:** Overview of ROSMAP single-cell RNA sequencing studies integrated in ROSMAP-Compass.

| Year | Author | Cohort | Synapse ID | Protocol |
| --- | --- | --- | --- | --- |
| 2019 | Mathys et al. | ROSMAP | syn3157322 | 10x |
| 2022 | Blanchard et al. | ROSMAP | syn2580853 | 10x |
| 2023 | Sun et al. | ROSMAP | syn51015750 | 10x |
| 2023 | Mathys et al. | ROSMAP | syn2580853 | 10x |
| 2024 | Mathys et al. | ROSMAP | syn52293417 | 10x |
| 2024 | Fujita et al. | ROSMAP | syn31512863 | 10x |
| 2025 | De Jager Lab | ROSMAP/AMP-AD | syn52339332 | 10x multiome |
| 2025 | Lee et al. | NPS-AD | syn52160016 | 10x multiome |

**Figure 1.**
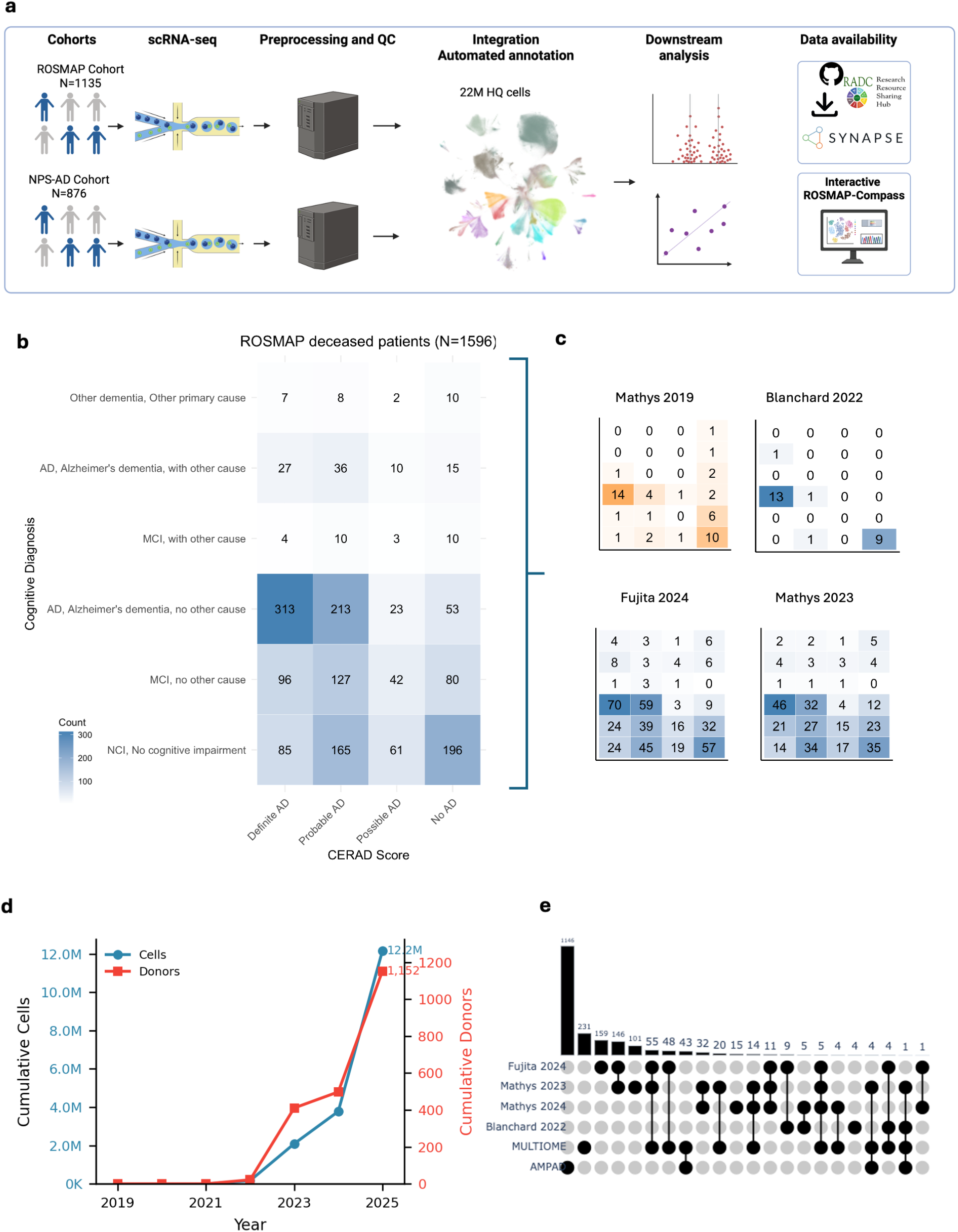
Overview of the ROSMAP single-cell brain atlas: study design, dataset expansion, and clinical characterization. **a**, Schematic workflow of the ROSMAP single-cell RNA sequencing study. Brain tissue samples from two cohorts (ROSMAP Cohort, N=1,135 and NPS-AD Cohort, N=876) were processed using 10x Genomics single-cell RNA sequencing. Following preprocessing and quality control, 22 million high-quality nuclei were integrated using automated annotation methods. The processed data underwent downstream analysis and are publicly available through multiple repositories including the AD Knowledge Portal, Synapse, and an interactive web-based visualization tool (ROSMAP-Compass). **b**, Heatmap showing the distribution of cognitive diagnoses across CERAD (Consortium to Establish a Registry for Alzheimer’s Disease) scores for deceased ROSMAP patients (N=1,596). Numbers in cells indicate patient counts, with darker blue shading indicating higher patient counts. **c**, heatmaps with same structure as in b, but for separate studies, orange indicates V2 chemistry was used, blue v3. **d**, Growth trajectory of the ROSMAP single-cell dataset from 2019 to 2025. The blue line indicates cumulative cell count (left y-axis) and the red line shows cumulative donor count (right y-axis). The dataset has expanded from initial pilot studies to over 12 million cells from more than 1,200 donors. **e**, UpSet plot comparing donor overlap across published single-cell RNA sequencing studies. Horizontal bars indicate the total number of donors per study (set size), with studies ordered by publication year. Vertical bars represent the number of donors shared between study combinations (intersection size), with connected dots below indicating which studies contribute to each intersection. Black dots and bars indicate unique or shared donors across the five studies (Fujita 2024, Mathys 2023, Mathys 2024, Blanchard 2022, MULTIOME, and AMPAD).

## 2 Results

To create ROSMAP-Compass, we systematically integrated all publicly available ROSMAP single-cell RNA sequencing data generated across multiple studies from 2019 to 2024. The primary challenge was harmonizing data from different laboratories, sequencing chemistries (10x v2 and v3), and computational pipelines into a unified resource. This required addressing technical variations including chemistry-specific biases, sample redundancies across studies, and inconsistent cell type annotations. Here, we describe the data landscape, the technical challenges we addressed during harmonization, and the architecture of the resulting integrated resource.

### 2.1 Overview of ROSMAP Single-Cell Sequencing identifies samples sequenced up to 4 times

We identified that as of now (August 2025), a total of 1,135 of ROSMAP patients has been sequenced using single-cell RNA sequencing technologies (Figure 1a). Notably, multiple donors have been sequenced repeatably using different chemistries and by different laboratories. Of note, 88 donors were sequenced 3 times and 6 donors sequenced even four times (Figure 1e). These samples provide an opportunity to compare sequencing results but at the same time it is essential to track them. Our analysis reveals that early single-cell studies did not necessarily reflect the full spectrum of AD pathology present in the ROSMAP cohort. We attribute this primarily to the limitations of early phase scRNA-seq technology, including high costs and the experimental nature of pilot projects during the initial adoption of single-cell methods (Mathys et al. 2019). This initial study was using 10x chromium V2 chemistry which is limited in compatibility with the later introduced 10x chromium V3 chemestry [1]. Despite these early limitations, the field reached a significant milestone in 2023 with the first major high-throughput study, encompassing 2.3 million cells from 400 donors. This substantial increase in scale enabled better statistical power and more accurately represents the diversity of the ROSMAP cohort (Mathys 2023, Figure 1bc). Integrating the most early and later studies poses another potential source of bias to account for in our large-scale integration effort.

### 2.2 Architecture of the Integrated ROSMAP Single-Cell Resource

Our integrated resource represents a substantial expansion beyond previous ROSMAP singlecell datasets. While earlier iterations included 8 million cells aligned using T2T methodology (including v2 chemistry data) with annotations based on the Siletti et al. [19] 3 million cell reference, we recognized limitations in both the chemistry and annotation framework. The current resource addresses these limitations through several key improvements. First, we excluded all v2 chemistry data to ensure consistent data quality. Second, we expanded the dataset to include additional data from the Mount Sinai NPS-AD study, resulting in 22 million cells that passed quality control metrics (Figure 2a,b.c). Of these, we successfully mapped 17 million cells to specific donors. We retained the 5 million unmapped cells in the dataset as they may provide valuable input for unsupervised model training, though they did not affect our downstream analyses as unlabeled cells were omitted from supervised approaches. We adopted the SEA-AD[10] reference annotation derived from the Siletti et al. study for this expanded dataset. This annotation framework aligns with ongoing efforts at the Allen Institute for Brain Science and facilitates seamless integration and comparison with other brain atlas initiatives. The dataset encompasses 24 broad cell types and 139 fine cell types, providing granular resolution for cell type-specific analyses. Demographically, while the dataset contains a higher absolute number of cells from male patients, more female participants have been sequenced overall, reflecting sampling depth variations across donors. The age distribution spans a remarkable range, from early childhood to elderly patients over 110 years of age, capturing the full spectrum of aging and age-related pathology. The integrated datasets show consistent clustering patterns across studies, demonstrating successful batch correction and harmonization (Supplemental Figure 2). The resource shows consistent data quality and marker gene expression across all component datasets (Supplemental Figure 3). Moreover, it captures the full demographic and pathological diversity of the ROSMAP cohort, with representation across age groups, Braak stages, CERAD scores, and APOE genotypes (Supplemental Figure 4).

**Figure 2.**
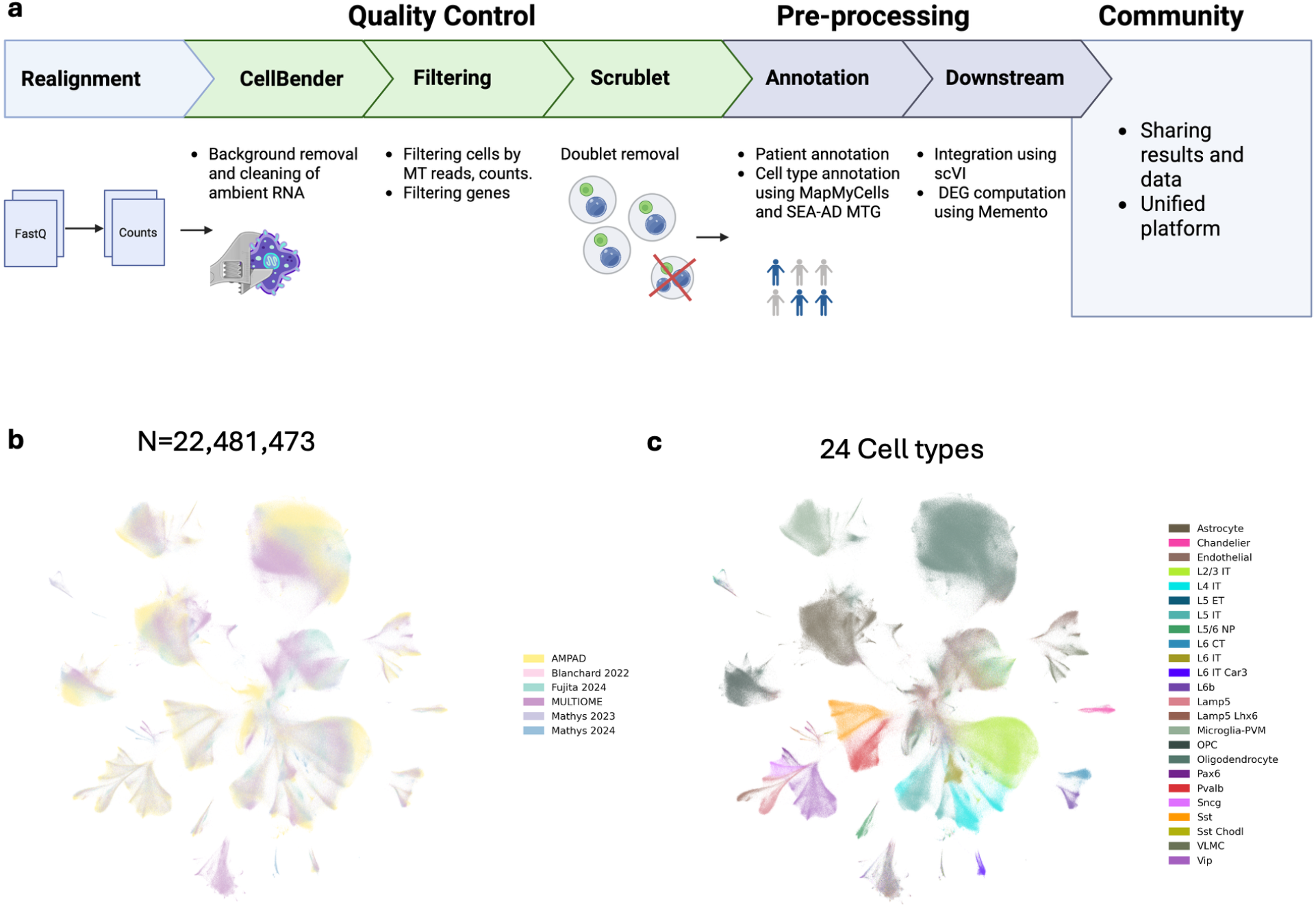
Single-nucleus RNA sequencing data processing and cellular diversity in the ROSMAP cohort. **a**, Schematic overview of the computational pipeline for single-nucleus RNA sequencing data analysis. The workflow comprises three main stages: quality control (including background removal with CellBender and filtering by mitochondrial reads, counts, and genes), pre-processing (doublet removal with Scrublet, patient annotation, and cell type annotation using MapMyCells and SEA-AD MTG), and community integration (using scVI and DEG computation with Memento). Data sharing is facilitated through a unified platform. **b**, UMAP visualization of all 22,481,473 nuclei colored by processing batch, demonstrating successful batch integration across the studies. **c**, UMAP visualization of the same nuclei colored by cell type identity, revealing 24 distinct cell types including astrocytes, oligodendrocytes, endothelial cells, various neuronal subtypes.

### 2.3 Democratizing Access to the Integrated ROSMAP Compass

To maximize the scientific impact of this integrated resource, we have implemented a multitiered access strategy that balances patient privacy requirements with research accessibility. The fully processed dataset is available through Synapse, adhering to the established data governance framework for the Rush Alzheimer’s Disease Center (RADC). While the RADC’s rigorous protocols for protecting patient rights and maintaining anonymity necessitate administrative procedures for data access, we emphasize that these safeguards represent best practices in human subjects research. The scientific value of this resource far exceeds the modest administrative investment required for access authorization.

Recognizing that not all researchers have the resources or time to navigate formal data access procedures, we have developed a complementary solution: a publicly available web server that provides immediate access to comprehensive analyses as they become available. This platform hosts all current analytical results and will be continuously updated with future analyses as they become available. The server includes extensive exploration of meta-analysis from both ROSMAP and NPS-AD cohorts, enabling researchers to interrogate the biological and clinical richness of these datasets without requiring local computational infrastructure, data application or registration. Our web platform will encompass both legacy analyses performed on T2T-aligned data and our primary hg38-aligned dataset, with ongoing development focused on the latter. This dual approach ensures that researchers can access immediate insights while pursuing formal data access for custom analyses, thereby maximizing the utility of this exceptional resource for the broader Alzheimer’s disease research community. ROSMAP-Compass includes differential expression analysis capabilities that enable researchers to explore gene expression patterns across clinical and demographic variables. Given the scale of possible comparisons when examining multiple conditions, sex-stratified analyses, and disease stages, we position ROSMAP-Compass as a hypothesis-driven resource optimized for targeted queries rather than genome-wide discovery. The web interface provides access to pre-computed analyses without multiple testing correction, allowing researchers maximum flexibility in their investigations. We recommend users approach the resource with specific hypotheses, searching for genes or pathways of interest based on prior knowledge. For rigorous statistical analysis, researchers should apply appropriate multiple testing corrections within their specific gene sets of interest. The resource is particularly valuable for validation of findings from other datasets, hypothesis testing for specific genes or pathways, candidate gene prioritization for follow-up studies, and cross-cohort comparisons with other AD datasets. For users requiring genome-wide discovery analyses, we recommend downloading the raw data to implement custom analysis pipelines with study-appropriate statistical corrections. This approach balances accessibility with statistical rigor, enabling researchers to leverage the full power of ROSMAP-Compass while maintaining appropriate analytical standards.

## 3 Conclusion and Future Work

We present ROSMAP-Compass: a fully harmonized and unified resource encompassing the entirety of single-cell RNA sequencing data from the ROSMAP cohort with comprehensive metadata integration. Through systematic re-alignment and curation, we have created a dataset of 22 million high-quality nuclei from 2,058 donors that addresses critical technical challenges including cross-study redundancies, chemistry-specific biases, and sample overlaps that previously complicated integrative analyses [7]. The resource is freely accessible through multiple tiers: immediate exploration via our web portal, full dataset access through Synapse following data use agreements, and upcoming natural language query capabilities through Model Context Protocol (MCP) integration [12, 8]. This tiered approach balances patient privacy requirements with scientific accessibility, enabling researchers regardless of computational resources to leverage this valuable dataset. Future development will focus on incorporating additional data modalities from ROSMAP including spatial transcriptomics and multiome data, expanding the MCP interface for enhanced natural language querying, and integrating new ROSMAP datasets as they become available. We will also maintain synchronization with evolving cell type annotations from brain atlas initiatives. We are committed to maintaining ROSMAP-Compass as a living resource that grows with the ROSMAP cohort.

## 4 Methods

### 4.1 Data Collection

We collected and harmonized single-cell RNA sequencing data from Alzheimer’s disease (AD) and control (CT) samples from the Religious Orders Study and Memory and Aging Project (ROSMAP)[4]. Specifically, we integrated data from four state-of-the-art studies as detailed in Supplementary Table 1. While initial assessment suggested only partial overlap between datasets (particularly between Mathys 2023 and Mathys 2024), we identified additional overlap with Blanchard et al. (2022), which was subsequently confirmed by the authors [5]. Notably, we discovered substantial overlap of 8 patients between Blanchard et al. (2022) and Mathys et al. (2019) comprising 10x Chromium v2 samples, also confirmed through author correspondence. All data were realigned directly from raw sequencing reads deposited on Synapse. This included one additional donor deposited under Synapse ID syn2580853 but not included in the Blanchard et al. (2022) publication, resulting in 25 unique donors from this dataset. After removing all redundancies and overlaps, our cleaned dataset comprises 1135 unique donors from the ROSMAP cohort across the four studies.

### 4.2 Data Processing V1: T2T Alignment

We acquired all metadata using the Synapse Python client (synapseclient v4.5.1) with Python 3.11.10. Count matrices were generated through realignment using Cell Ranger v8.0.1 [22]with the T2T reference genome [18]. To ensure data integrity, we systematically removed redundancies by identifying overlapping cells and shared donors across studies, excluding all duplicated cells from downstream analyses. Quality control thresholds were applied as follows: genes were required to be expressed in at least 3 cells, and cells expressing fewer than 100 unique genes were discarded. Putative doublets were removed using Scrublet v0.2.2 [21]) with default parameters. Ambient RNA correction was performed using CellBender v0.3.2 [6] with default settings. Cell type annotation was performed using the Allen Brain Institute’s MapMyCells tool [2]. Only cell clusters containing more than 100 cells were retained for subsequent analyses.

### 4.3 Data Processing V2: GRCh38 Alignment

The second iteration of our processing pipeline followed identical quality control procedures but utilized the GRCh38 reference genome. We employed the same software versions: Synapse Python client (v4.5.1) with Python 3.11.10 and Cell Ranger v8.0.1. Redundancy removal, quality control thresholds, doublet detection with Scrublet v0.2.2, and ambient RNA correction with CellBender v0.3.2 were applied consistently with the T2T pipeline.

### 4.4 Differential Gene Expression Analysis

Differential gene expression between AD and control groups was computed using MEMENTO [13] for each sex separately computed across all cell types and conditions.

## 5 Code availability

All computed DEGs will be available on the official ROSMAP-Compass website upon publication. All code will be available on GitHub https://github.com/ROSMAP-Compass this will also serve to update about coming updates and websites.

## 6 Data availability

All data is freely available on synapse.org syn2580853, syn2580853, syn52293417, syn2580853, syn52339332. The processed atlas is available at syn65460889.

## 7 Author contributions

Conceptualization & design: M.F. Writing – original draft: M.F. Implementation: M.F., I.F.D. Website implementation: M.F, I.F.D. Technical expertise & methodology: M.F., I.F.D., P.F., P.H. F.G. A.K. Resources & funding acquisition: A.K. All authors contributed to discussions, provided critical feedback, and approved the final manuscript.

## 8 Supplementary Information

**Supplementary figure 1.**
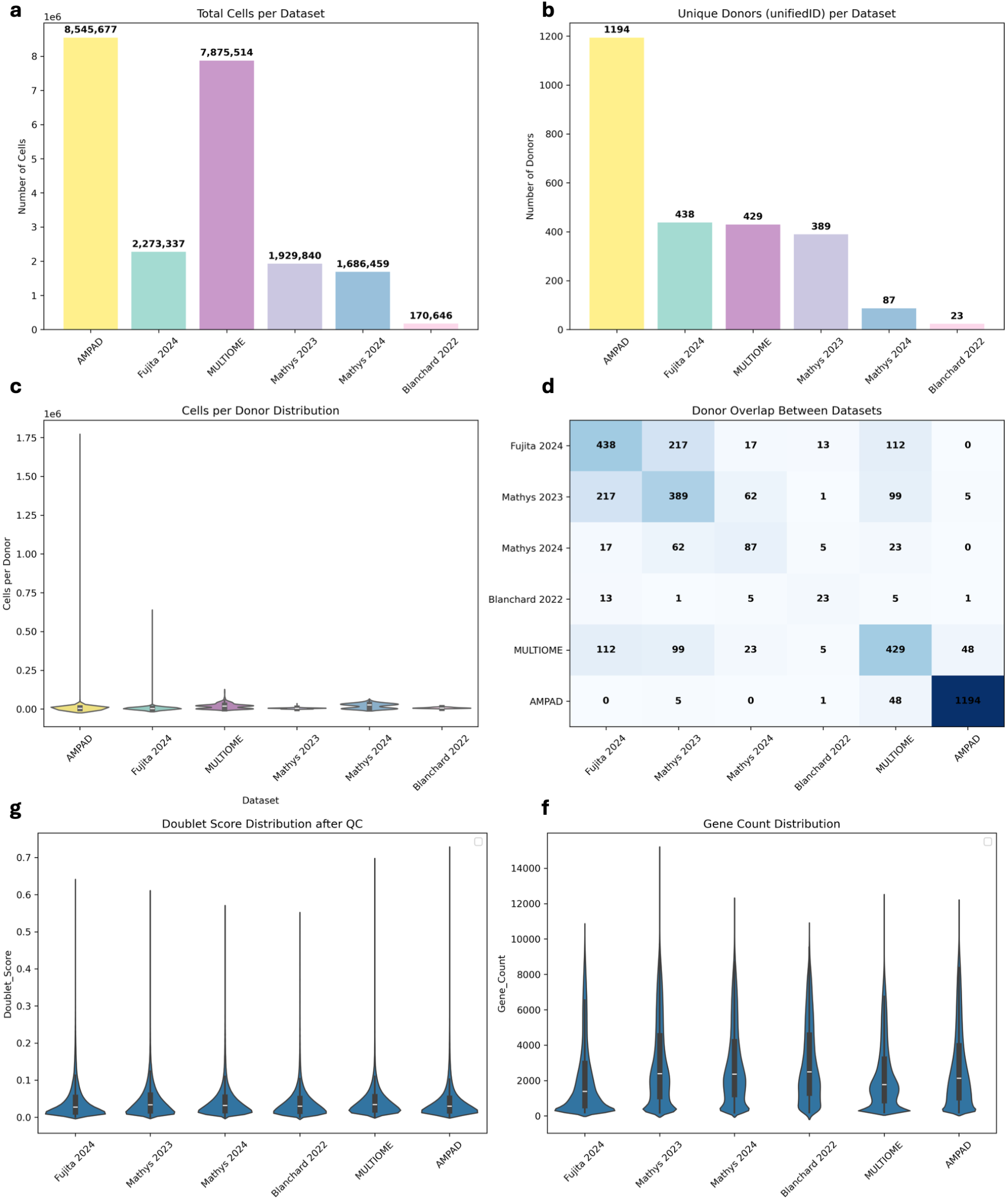
**a**, Total cell counts per dataset showing variation in sample size across studies. **b**, Number of unique donors per dataset after unifiedID assignment. **c**, Violin plots showing the distribution of cells per donor across datasets, with median values indicated. **d**, Heatmap displaying donor overlap between dataset pairs, with numbers indicating shared donors. **e**, Doublet score distribution after quality control filtering, demonstrating consistent low doublet rates across all datasets. **f**, Gene count distribution per cell across datasets, showing comparable sequencing depth and data quality.

**Supplementary figure 2.**
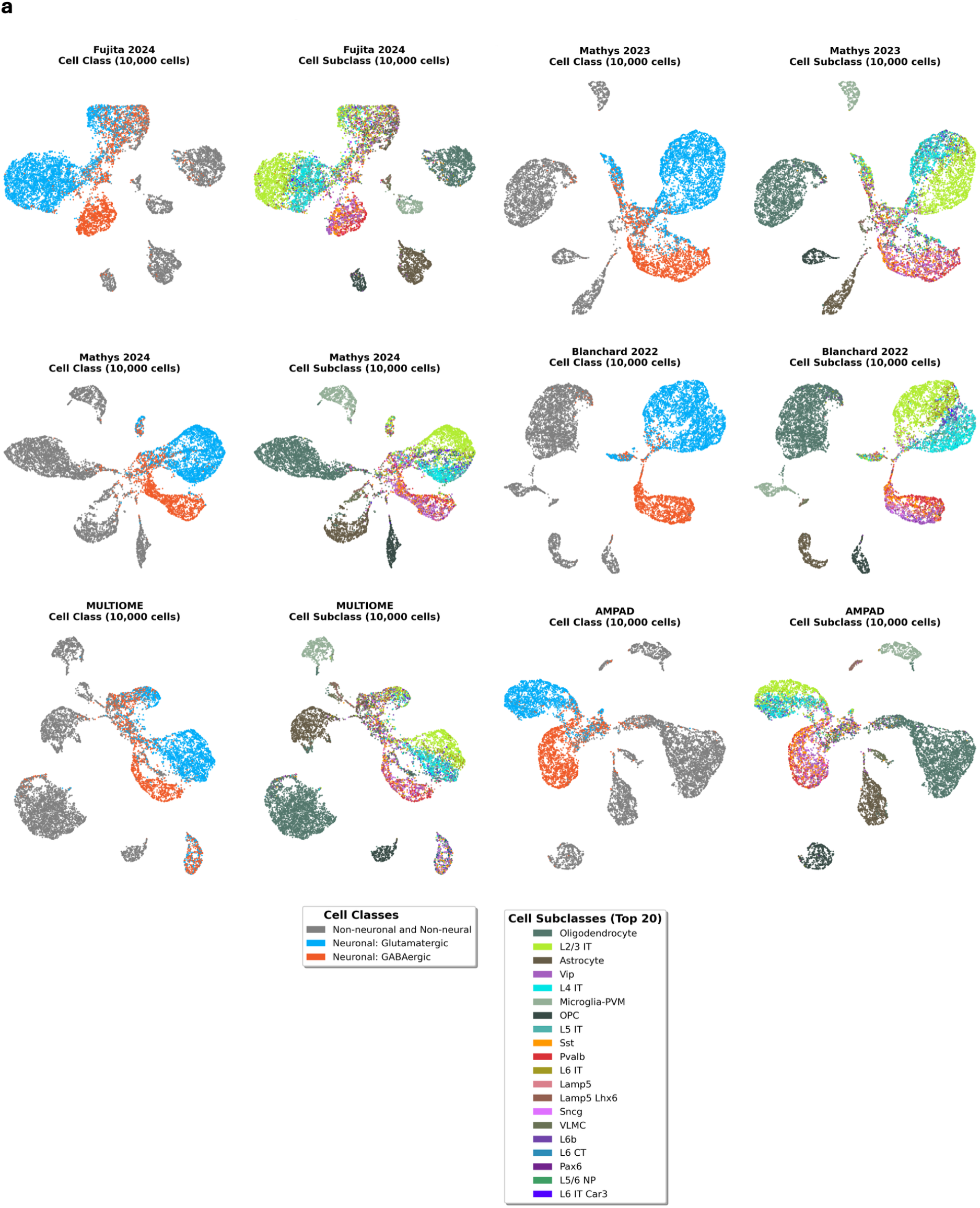
**a**, UMAP projections of integrated single-cell RNA-seq data from six datasets (Fujita 2024, Mathys 2023, Mathys 2024, Blanchard 2022, MULTIOME, and AMPAD), each subsampled to 10,000 cells. Left panels show cells colored by major cell class (neuronal, glutamatergic, and GABAergic). Right panels display the same projections colored by cell subclass identity (top 20 subclasses shown), demonstrating consistent cell type clustering across datasets after integration.

**Supplementary figure 3.**
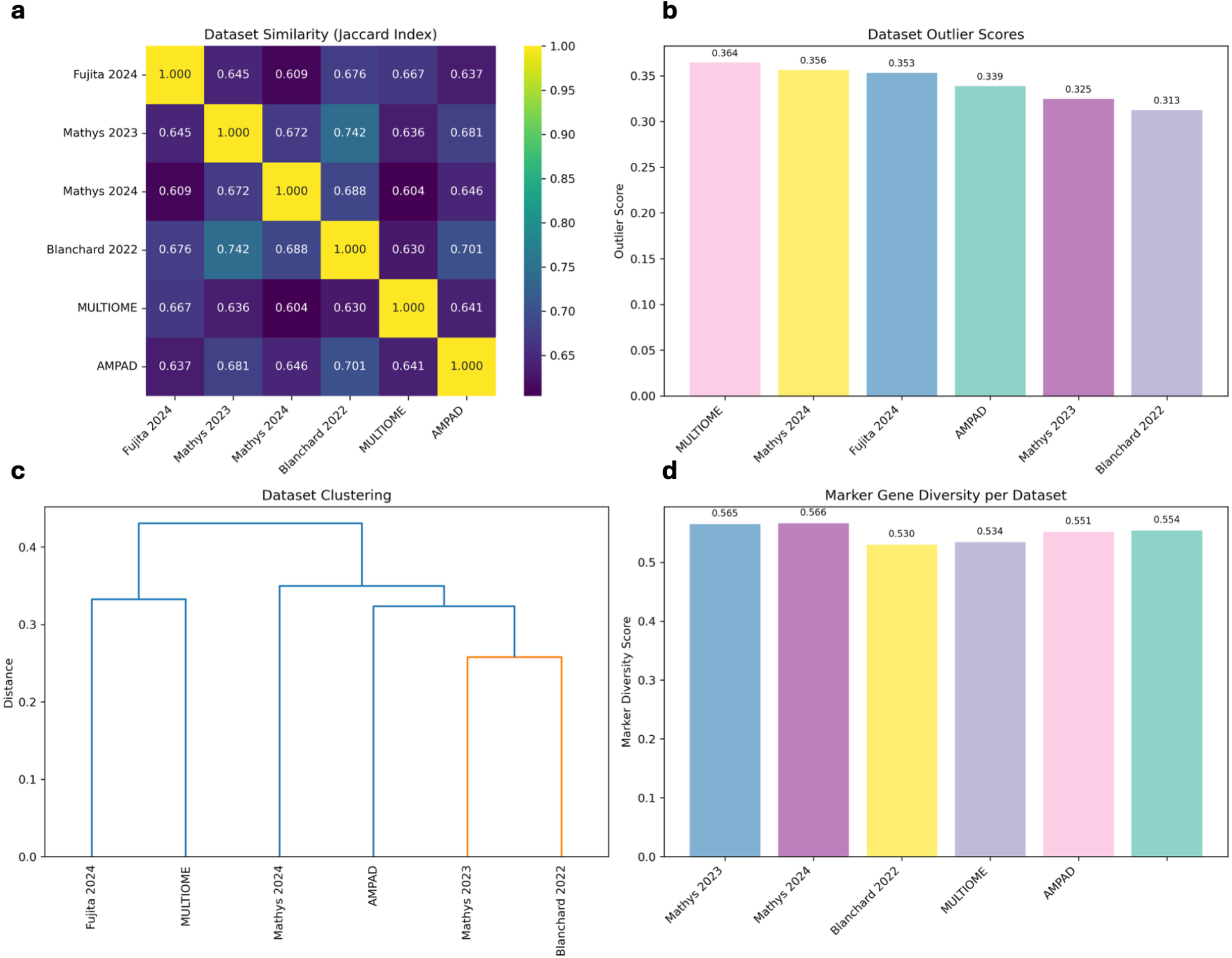
**a**, Heatmap showing pairwise dataset similarity based on Jaccard index, with values ranging from 0 (low similarity) to 1 (high similarity, diagonal). **b**, Bar plot of dataset outlier scores, indicating relative data quality across studies. **c**, Hierarchical clustering dendrogram based on dataset similarity, revealing relationships between studies. **d**, Marker gene diversity scores per dataset, demonstrating consistent capture of cell type-specific gene expression across all studies.

**Supplementary figure 4.**
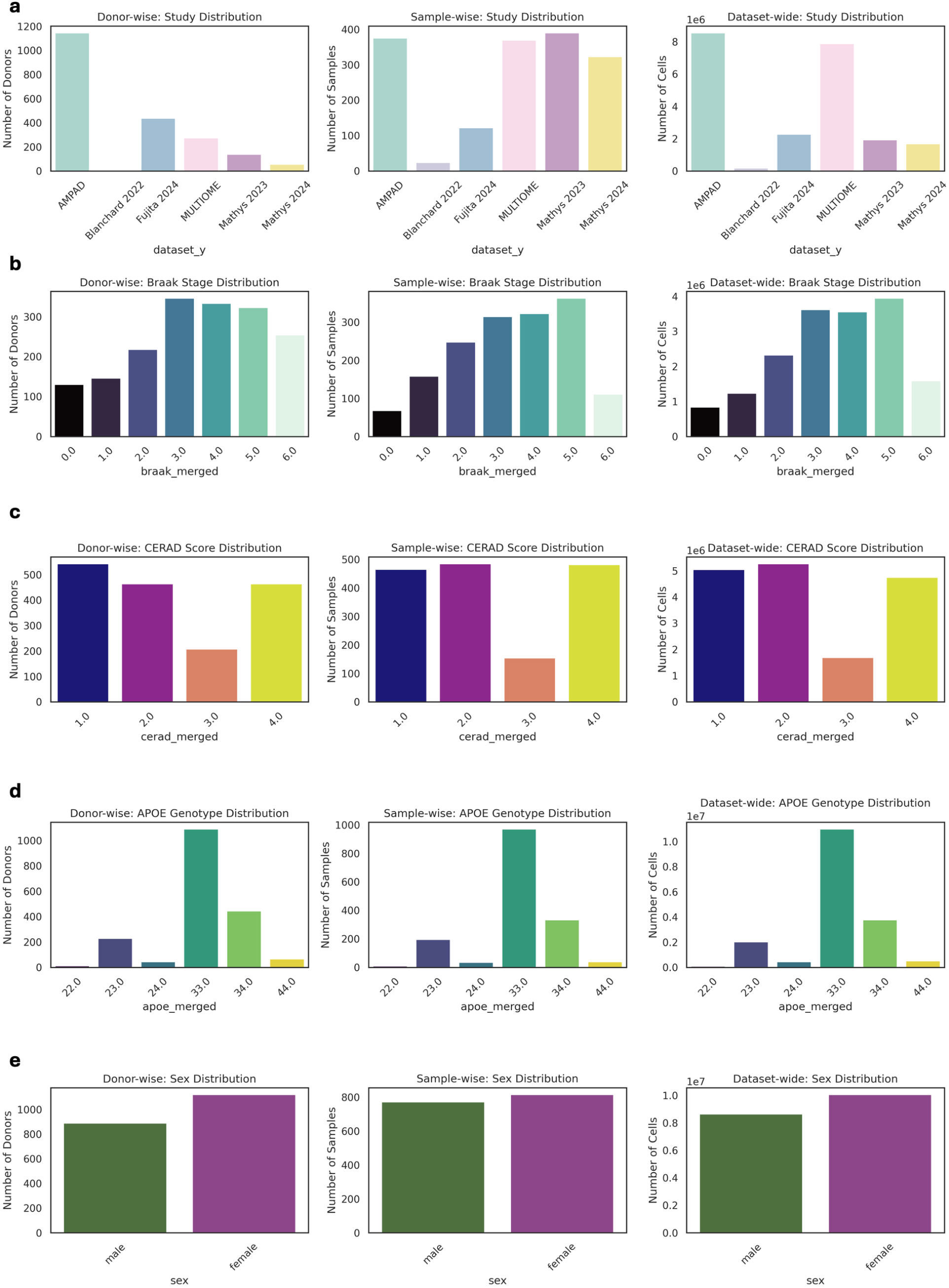
**a**, Distribution of studies across donor-wise, sample-wise, and dataset-wise levels, showing representation of each dataset in the integrated analysis. **b**, Braak stage distribution at donor, sample, and dataset levels, demonstrating coverage of Alzheimer’s disease pathological stages (0-6.0) across the cohort. **c**, CERAD score distribution showing neuropathological plaque density assessments (1.0-4.0) at donor, sample, and dataset levels. **d**, APOE genotype distribution displaying frequency of APOE*ε*2/3, APOE*ε*3/3, APOE*ε*3/4, and APOE*ε*4/4 alleles across donor, sample, and dataset levels. **e**, Sex distribution showing male and female representation at donor, sample, and dataset levels across the integrated cohort.

## 9 Competing interests

No competing interest is declared.

## 10 Author contributions statement

Conceptualization & design: M.F. Writing – original draft: M.F. Implementation: M.F., I.F.D. Website implementation: I.F.D., M.F. Biological expertise & methodology: M.F., I.F.D., V.W., P.H., F.G., P.F., A.K.

## 11 Acknowledgments

This study was financed through the DFG project 469073465, and the M.J. Fox Foundation (MJFF-021418; A.K. & T.W-C.). Figures were created with BioRender.com.

## Notes

https://github.com/ROSMAP-Compass

### Competing Interest Statement

The authors have declared no competing interest.

